# Pan-Cancer Exploration of mRNA Mediated Dysregulated Pathways in the Cancer Genomics Cloud

**DOI:** 10.1101/599225

**Authors:** Margaret Linan, Junwen Wang, Valentin Dinu

## Abstract

We performed a comprehensive pan-cancer analysis in the Cancer Genomics Cloud of HTSeq-FPKM normalized protein coding mRNA data from 17 cancer projects in the Cancer Genome Atlas, these are Adrenal Gland, Bile Duct, Bladder, Brain, Breast, Cervix, Colorectal, Esophagus, Head and Neck, Kidney, Liver, Lung, Pancreas, Prostate, Stomach, Thyroid and Uterus. The PoTRA algorithm was applied to the normalized mRNA protein coding data and detected dysregulated pathways that can be implicated in the pathogenesis of these cancers. Then the PageRank algorithm was applied to the PoTRA results to find the most influential dysregulated pathways among all 17 cancer types. Pathways in cancer is the most common dysregulated pathway, and the MAPK signaling pathway is the most influential (PageRank score = 0.2034) while the purine metabolism pathway is the most significantly dysregulated metabolic pathway.

## INTRODUCTION

Recent studies have shown that both dysregulated or frequently mutated pathways should be used for the characterization of cancers instead of driver mutations (Zhang, Chien, Yong and Kuang, 2017). Network biology is an important approach that can detect frequently dysregulated pathways in distinct cancer types (Zhang, Chien, Yong and Kuang, 2017). Dysregulated pathways are biological networks that have collections of hub genes that are significantly different between cancer and normal tissues. Hub genes can act individually to impact the function of other genes or entire biological networks (Flintoft, 2004). Functional alterations in tumors have also been found to cause a cascade of alterations in pathway networks (Sanchez-Vega et al., 2018). These alterations target pathways that are advantageous for the tumors and thus avoid targeting pathways that may lead to cellular death (Sanchez-Vega et al., 2018). Identifying mRNA mediated dysregulated pathways through the analysis of large scale gene expression data has been done with TCGA data before but only with smaller numbers of cancer types that did not further consider the subtypes of these cancers or their intra-tumoral variation. In the present work, the Pathways of Topological Rank Analysis (PoTRA) algorithm was used to build correlation based gene networks from TCGA open-access data for 17 cancer types. Specifically, the PoTRA algorithm finds the correlation between each phenotypes gene expression set and the genes associated with each KEGG pathway then constructs networks for each (Li, Liu and Dinu, 2018). Next, the dysregulated KEGG pathways with topologically ranked hub genes that are significantly different between normal and cancer are detected using Fisher’s exact test (Li, Liu and Dinu, 2018). Additionally, dysregulated KEGG pathways with significantly different distributions of PageRank scores of topologically ranked genes are also detected using the Kolmogorov–Smirnov test (Li, Liu and Dinu, 2018). The PoTRA algorithm focuses on protein-coding messenger RNA (mRNA) mediated dysregulated pathways because mRNAs are part of the stress response at the translational level and also because they are mediators of carcinogenesis (Vaklavas, Blume and Grizzle, 2017).

Other network based methods for detecting dysregulated pathways includes multi-omics approaches where both gene mutations with high coverage and no gene overlap (single nucleotide variants, copy numbers) and gene expression data are analyzed with the Dendrix and Markov Chain Monte Carlo algorithms (Wu, Dong and Wei, 2017). Additional approaches include using subnetworks from pathway interaction networks to detect dysregulated pathways by treating the detection as a feature selection task (Liu, Liu, Hao, Chen and Zhao, 2012). Another approach uses a combination of methylation and gene expression data with an autoencoder to identify dysregulated pathways by using the differential expression profiles of select genes (Visakh and Nazeer, 2018).

## MATERIALS AND METHODS

Google Cloud and Docker were utilized in the creation of containers for multiple data management and analysis algorithms including PoTRA. Rabix composer was utilized to match these containers with their corresponding command lines and to port the applications to the Seven Bridges Genomics Cancer Genomics Cloud (Lau et al, 2017) where they were implemented. The applications were applied to TCGA transcriptome profiling gene expression quantification HTSeq-FPKM normalized samples (normal and tumor) from 17 primary sites. These are Adrenal Gland, Bile Duct, Bladder, Brain, Breast, Cervix, Colorectal, Esophagus, Head and Neck, Kidney, Liver, Lung, Pancreas, Prostate, Stomach, Thyroid and Uterus.

The pre-processing algorithms removes the last few lines of statistical metrics from each sample, adds a header and then comma delimits the data. The purpose of the inner join stage is to join all of the tumor and normal data by their ENSG ID. The gene conversion stage converts the ENSG ID column in the joined data set to Entrez IDs. The random resampling stage creates new balanced data sets with equal numbers of tumor and normal samples. The pre-processing pipeline code is provided as supplemental information. The PoTRA algorithm “PoTRA_CorN” applies Fisher’s exact and Kolmogorov-Smirnoff tests as well as calculates the number of hubs and edges to support the detection of dysregulated pathways.

It also applies the PageRank algorithm (Page et al., 1999) to determine the topological importance of each pathway gene for both normal and tumor samples. Finally, it ranks the pathways and arrives at an average rank, then averages the metrics for the hubs and edges for normal and tumor samples. The post-processing aggregation algorithm applies the log sum function to p-values from the Fisher’s exact and Kolmogorov-Smirnoff tests. The list of significantly dysregulated pathways (Fisher’s Exact P-values < 0.05) from the 17 cancer projects were binned into six categories: Top 10, Top 20, Top 30, Top 40, Top 50, Top 60 then visualized in Neo4J, where they were further analyzed with the PageRank algorithm. Principal components analysis was performed to determine the distribution of normal and tumor for each primary site.

Different combinations of normal and tumor were tested to determine the optimal sample sizes for both phenotypes so that the average standard deviation in the ranks of the dysregulated pathways was minimized (Linan and Dinu, 2018).

The KEGG database was queried to determine how much of an overlap there is between the reported dysregulated pathways for each cancer type and the dysregulated pathways that were detected and ranked by PoTRA. A literature search was performed to validate the reported dysregulated pathways.

## RESULTS

### TCGA Samples

Only 17 primary sites of cancer had at least 3 samples for normal and tumor samples (Table 1). All of the HTSEQ FPKM normalized datasets that were created from the normal and tumor samples for each cancer type had more tumor samples than normal, and therefore were considered to be unbalanced.

**Table 1.** The sample sizes for each phenotype by primary site.

| Table 1 The sample sizes for each phenotype by primary site. |  |  |
| --- | --- | --- |
| Primary Site of Cancer | Normal Samples | Tumor Samples |
| Adrenal Gland | 3 | 257 |
| Bile Duct | 9 | 36 |
| Bladder | 19 | 414 |
| Brain | 5 | 667 |
| Breast | 113 | 1102 |
| Cervix | 3 | 304 |
| Colorectal | 51 | 644 |
| Esophagus | 11 | 161 |
| Head and Neck | 44 | 500 |
| Kidney | 128 | 891 |
| Liver | 50 | 371 |
| Lung | 108 | 1035 |
| Pancreas | 4 | 177 |
| Prostate | 52 | 498 |
| Stomach | 32 | 375 |
| Thyroid | 58 | 502 |
| Uterus | 35 | 607 |

### Cancer Subtypes

Only 6 of the 17 cancer types (Table 1) have subtypes that are available as TCGA projects.

1. Adrenal Gland: Adrenocortical Carcinoma (ACC), Pheochromocytoma and Paraganglioma (PCPG)
2. Brain: Glioblastoma Multiforme (GBM), Brain Lower Grade Glioma (LGG)
3. Colorectal: Colon Adenocarcinoma, Rectum Adenocarcinoma
4. Kidney: Kidney Renal Clear Cell Carcinoma (KIRC), Kidney Chromophobe (KICH), Kidney Renal Papillary Cell Carcinoma (KIRP)
5. Lung: Lung Adenocarcinoma (LUAD), Lung Squamous Cell Carcinoma (LUSC)
6. Uterus: Uterine Carcinosarcoma (UCS), Uterine Corpus Endometrial Carcinoma (UCEC)

Among the 13 cancer subtypes, (Figures 2-4) the subtypes where normal and tumor groups overlap are:

- ACC Tumor / PCPG Normal and Tumor
- COAD/READ Normal - COAD/READ Tumor
- GBM Normal - LGG Tumor
- KICH/KIRC Normal - KIRC Tumor
- KIRP Normal - KICH Tumor
- LUAD/LUSC Normal - LUAD/LUSC Tumor
- UCEC Normal - UCEC/UCS Tumor

**Figure 1.**
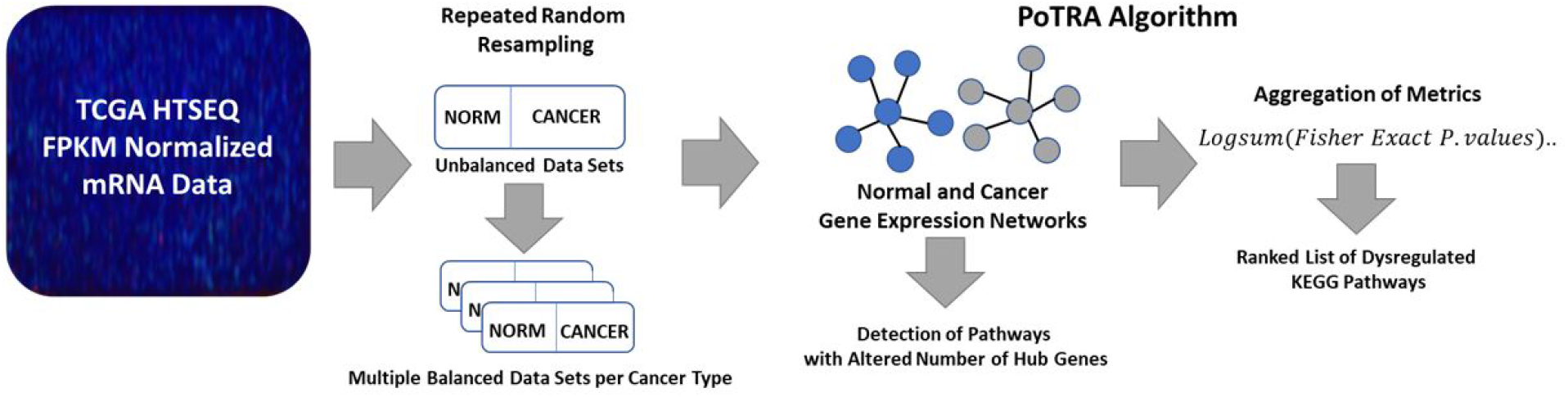
The custom CGC workflow for the detection of dysregulated pathways in TCGA HTSeq-FPKM normalized protein-coding mRNA normal and cancer samples. The mRNA data from cancer projects were retrieved from the CGCs TCGA portal and then pre-processed. The joined samples formed unbalanced data sets and therefore, underwent repeated random resampling. After the data sets were resampled, they formed multiple balanced data sets, that were then analyzed with the PoTRA algorithm. PoTRA detected several dysregaulted pathways per data set that were then aggregated and ranked with the aggregation algorithm.

**Figure 2.**
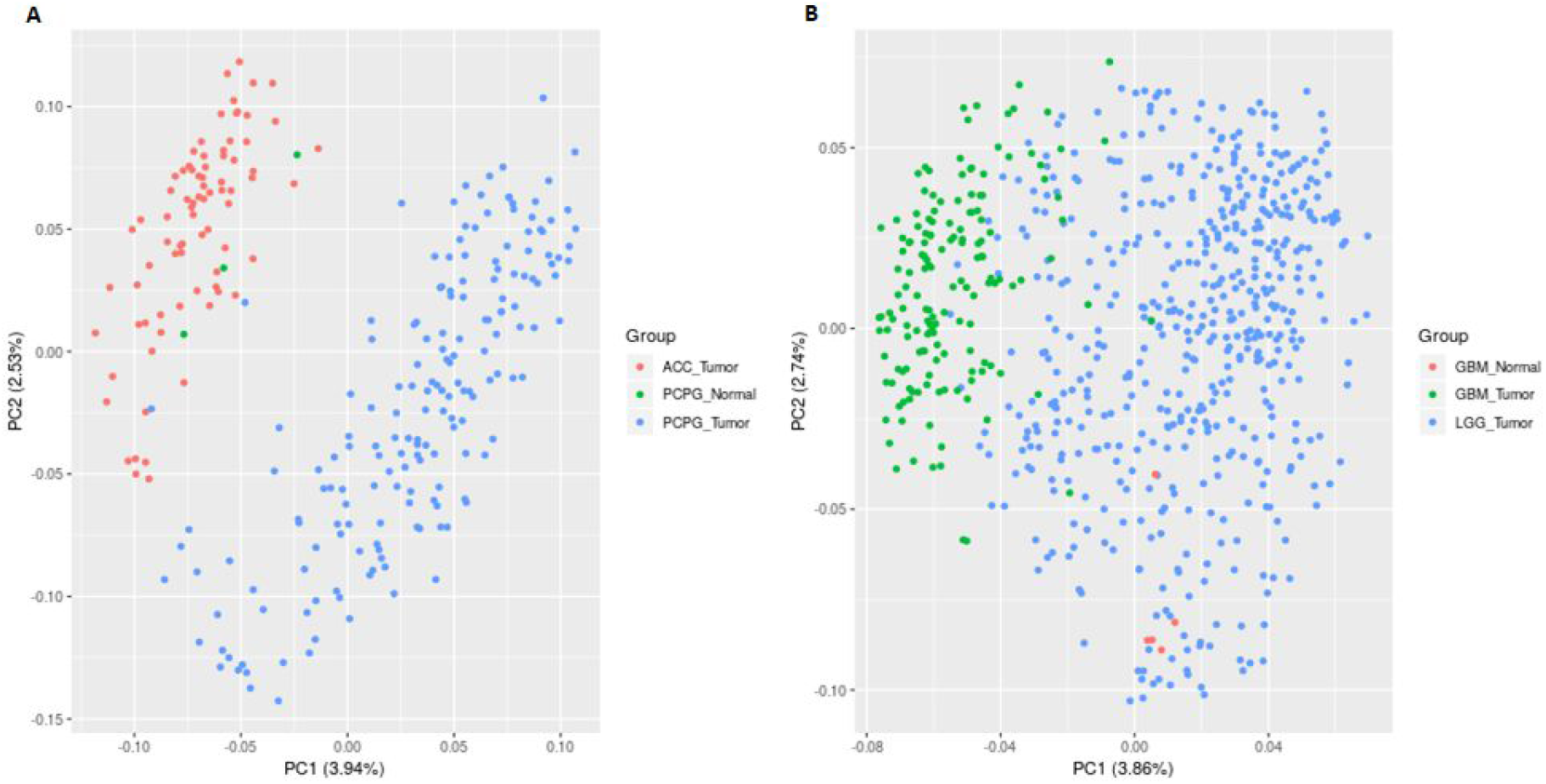
PCA plots of the normal and tumor HTSeq FPKM normalized miRNA data from the following TCGA projects a) ACC, PCPG, b) GBM and LGG. A) The ACC and PCPG project data have no overlap between their tumor gene expression values. Although only 3 samples were available for the PCPG normal group, this group overlaps with the expression values of the ACC tumor group. B) The GBM and LGG project data has a miminal overlap between the GBM and LGG tumor groups, there is also overlap between the GBM normal and LGG tumor groups gene expression values.

**Figure 3.**
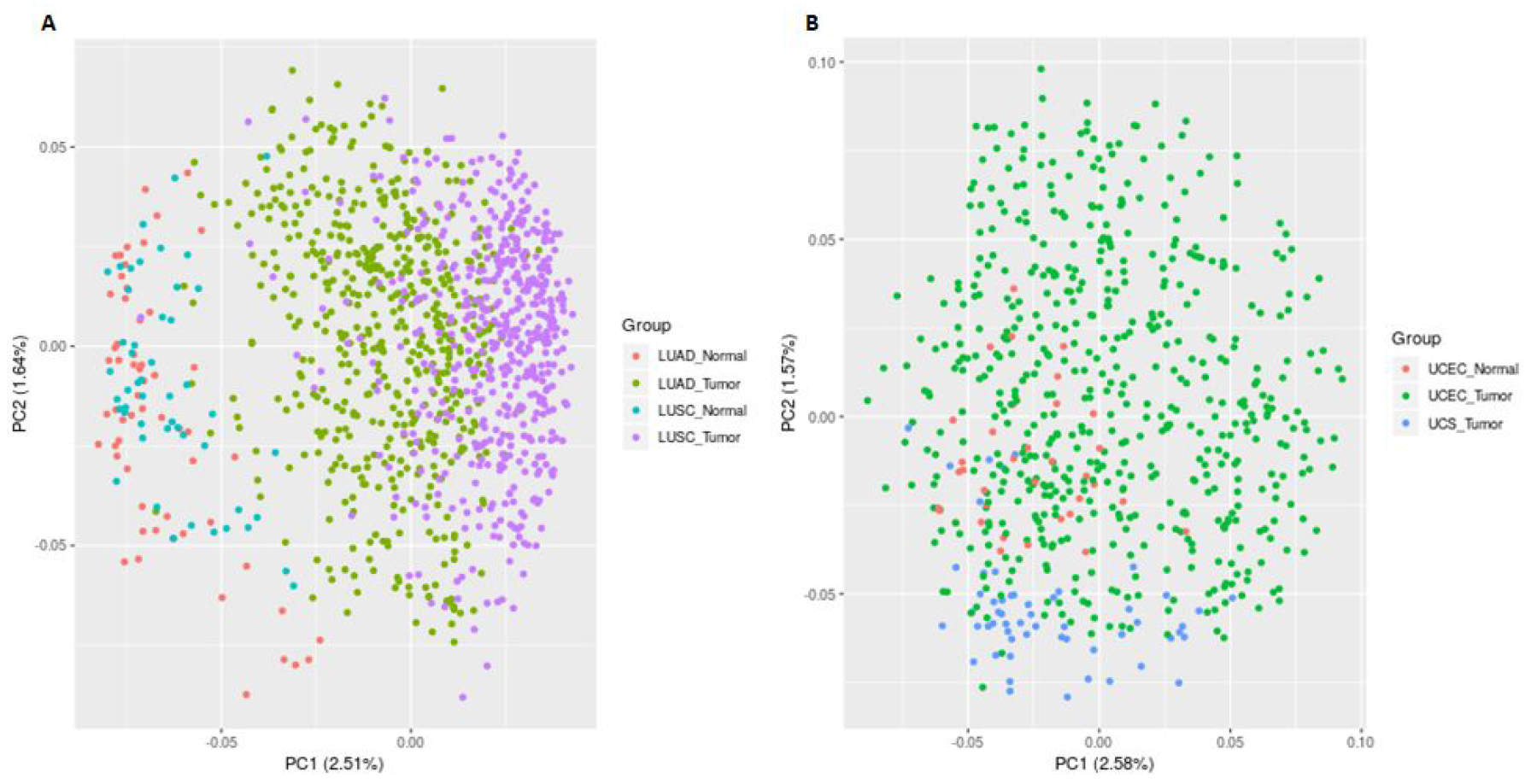
PCA plots of the normal and tumor HTSeq FPKM normalized miRNA data from the following TCGA projects a) LUAD, LUSC b) UCS and UCEC. A) The gene expression data from the LUAD and LUSC projects have overlap between their normal groups, and also have overlap between their tumor groups. B) In contrast, for the gene expression values of the UCS and UCEC projects, all of the normal and tumor groups overlap.

**Figure 4.**
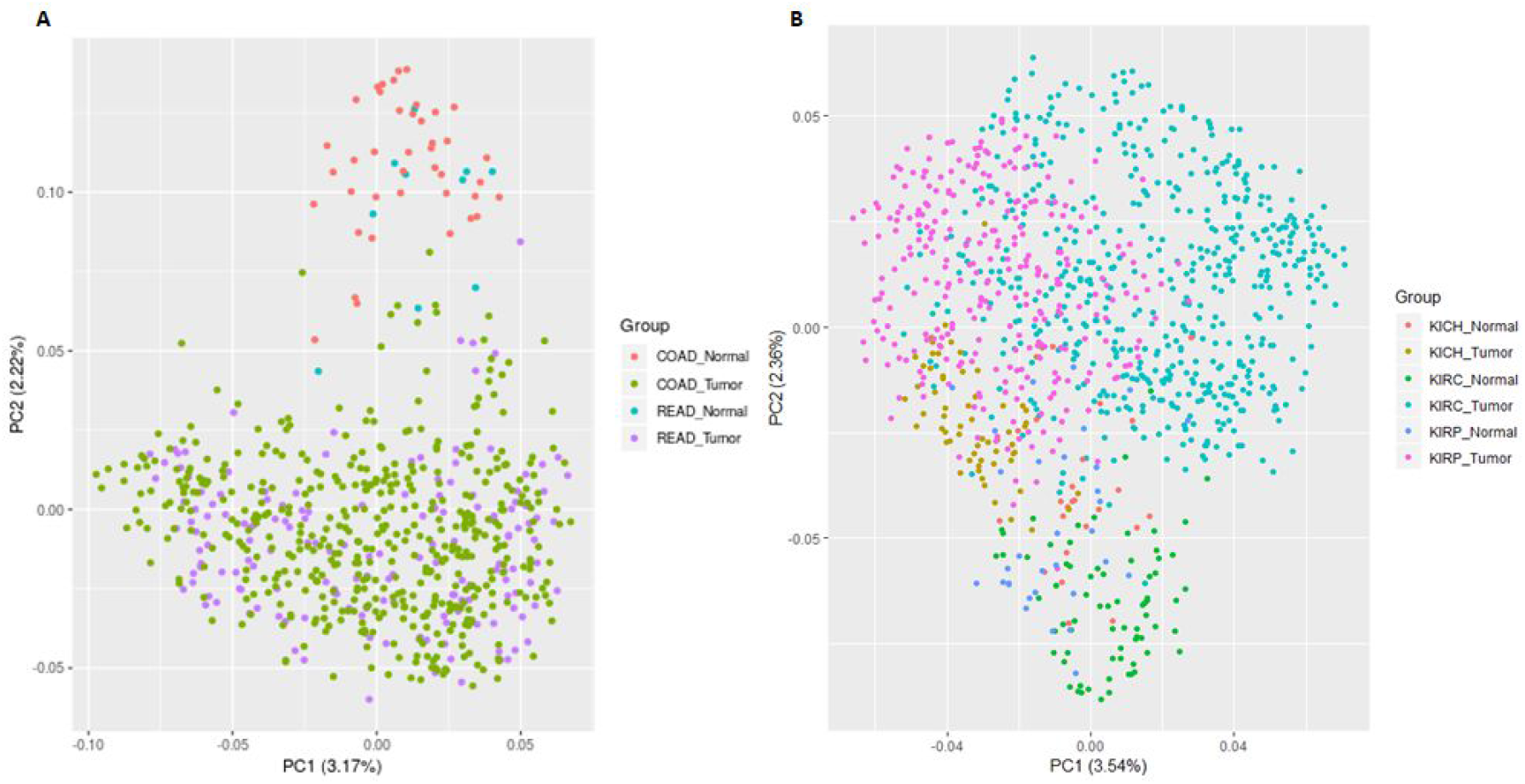
PCA plots of the normal and tumor HTSeq FPKM normalized miRNA data from the following TCGA projects a) COAD, READ b) KICH, KIRC and KIRP. A) The gene expression data from the COAD and READ projects have overlap between their normal groups, and also have overlap between their tumor groups. B) Similarly, for the gene expression values of the KICH. KIRC and KIRP projects, there are overlaps between their normal groups, and also between their tumor groups.

The subtypes with the most overlap between tumor groups are:

- COAD and READ
- KIRP, KIRC and KICH
- LUAD and LUSC
- UCS and UCEC

The subtypes with the least overlap between tumor groups are:

- ACC and PCPG
- GBM and LGG

### Dysregulated Pathways

The results of the pan-cancer analysis are a list of ranked mRNA mediated dysregulated pathways for 17 cancer types. Thus the results (Figure 5) can be interpreted as how many binned dysregulated pathways (logsum(Fisher’s Exact P-Value) < 0.05 and Rank >= 60) are associated with a cancer.

**Figure 5.**
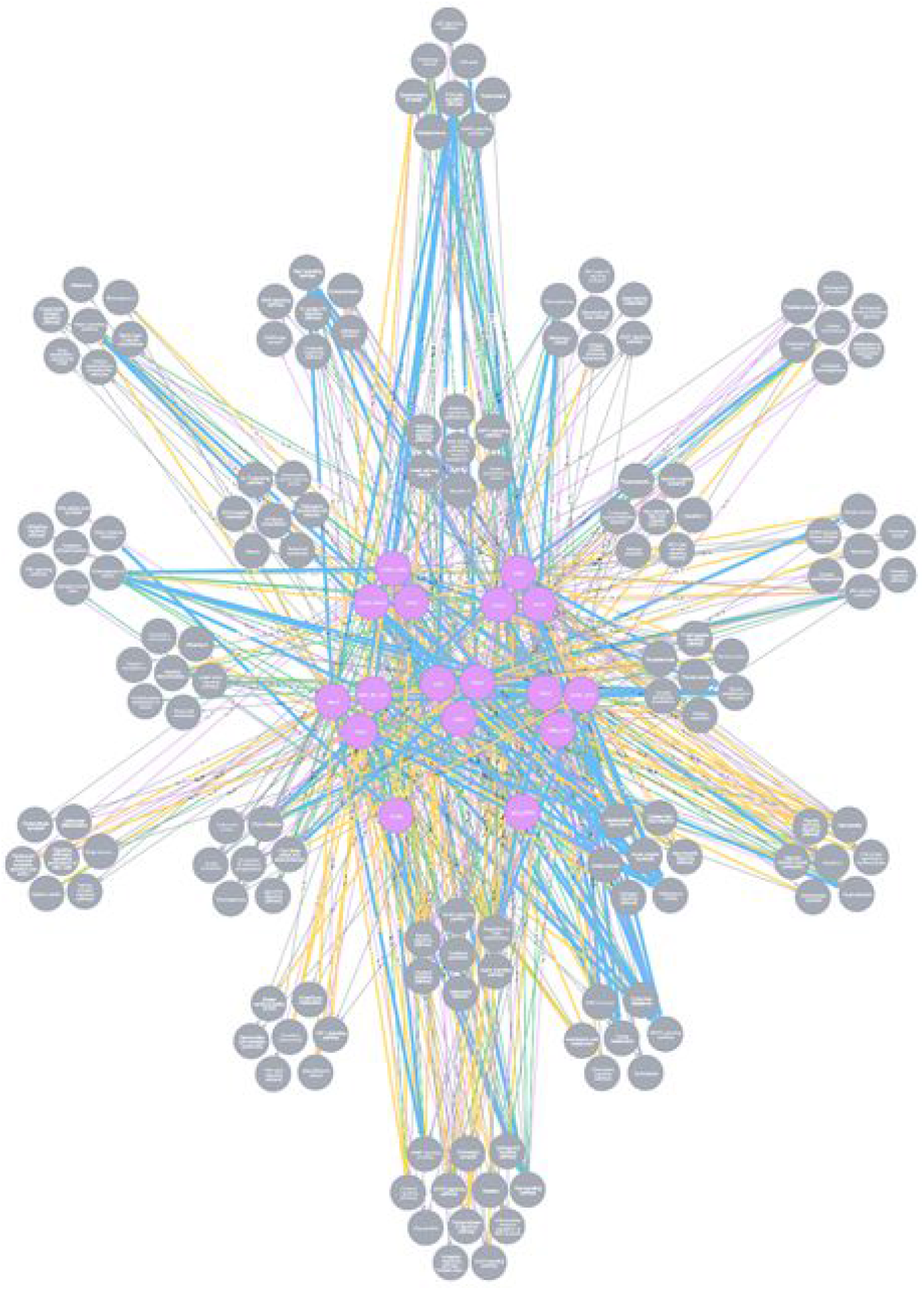
The pan-cancer graph network includes the names of the dysregulated pathways and the associated cancer projects from the TCGA(GRCh38 version). After PoTRA was used to detect dysregulated pathways (dark gray nodes) in 17 TCGA primary sites (faschia nodes), the Cypher language was used to build a graph network in Neo4J that represents the relationships between the dysregulated pathways and their associated cancer projects. The ranked pathways were binned into the following categories: Top 10 (blue). Top 20 (green). Top 30 (yellow). Top 40 (light purple). Top 50 (gray) and Top 60 (gray).

**Figure 6.**
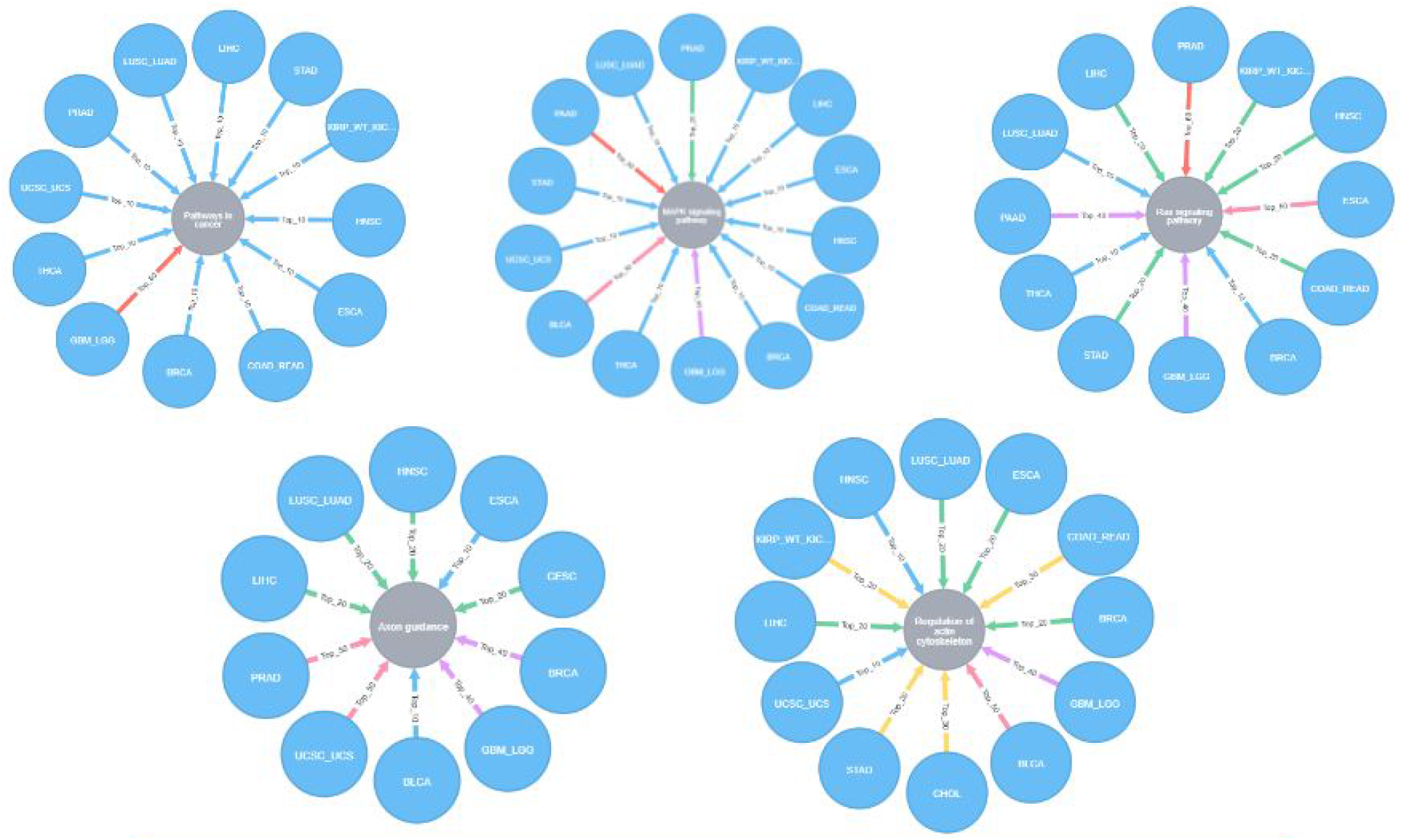
The top 5 dysregulated pathways that were detected by PoTRA. The graph networks depict the rankings of each pathway (Pathways in Cancer, MAPK signaling pathway. Ras signaling pathway, Axon guidance and Regulation of actin cytoskeleton) for several cancer projects. The Pathways in Cancer pathways is the most highly ranked dysregulated pathway. The MAPK signaling pathway is the 2nd most highly ranked dysregulated pathway, and Ras signaling pathway is the 3rd most highly ranked, while Regulation of action cytoskeleton is the 4th most highly ranked and Axon guidance is the 5th most highly ranked dysregulated pathway.

The Pathways in cancer, mitogen-activated protein kinase (MAPK), Ras signaling pathways as well as the Axon guidance and Regulation of actin cytoskeleton pathways were the most represented in the Top 10 category for most dysregulated pathways. In KEGG, the Pathways in cancer network contains 15 signaling pathways, that also includes the MAPK signaling pathway (Kanehisa, Furumichi & Tanabe et al., 2017; Kanehisa, Sato & Kawashima et al., 2016; Kanehisa & Goto, 2000). The MAPK signaling pathway is highly conserved and involved in the regulation of several cellular functions such as proliferation, differentiation, migration and transformation (Lake et al., 2016; Kanehisa, Furumichi & Tanabe et al., 2017; Kanehisa, Sato & Kawashima et al., 2016; Kanehisa & Goto, 2000; Slattery et al., 2018). Similarly, the Ras signaling pathway controls the regulation of cellular processes such as proliferation, differentiation, migration, survival and growth (Kanehisa, Furumichi & Tanabe et al., 2017; Kanehisa, Sato & Kawashima et al., 2016; Kanehisa & Goto, 2000). The Axon guidance pathway regulates the formation of the neuronal network and includes the MAPK signaling and regulation of actin cytoskeleton pathways in the axon repulsion process (Kanehisa, Furumichi & Tanabe et al., 2017; Kanehisa, Sato & Kawashima et al., 2016; Kanehisa & Goto, 2000). The Regulation of actin cytoskeleton pathway encompasses a number of processes that are vital for several cellular functions (Lee and Dominguez, 2010). The malfunction and disorganization of cytoskeletal proteins has been implicated in pathogenesis of many diseases and increased tumorigenicity (Segarra, Yavorski and Blanck, 2017).

The KEGG pathway known as Pathways in cancer, is the most common pathway in the Top 10 most dysregulated pathways in 11 of 17 TCGA cancer types. However, it was in the top 60 most dysregulated pathways for the Glioblastoma Multiforme and the Brain Lower Grade Glioma cancers. The MAPK signaling pathway is the most common (14 cancer projects / 17 total) and influential (PageRank score = 0.2034) among all of the significant topologically ranked dysregulated pathways for each of the 17 cancer projects (Table 2).

The MAPK signaling pathway is in the top 10 most dysregulated pathways list for the following cancers: Breast Invasive Carcinoma (BRCA), Esophageal Carcinoma (ESCA), Head and Neck Squamous Cell Carcinoma (HNSC), Liver Hepatocellular Carcinoma (LIHC), Colon and Rectum Carcinomas (COAD and READ), Kidney Renal Clear Cell Carcinoma (KIRC), Kidney Chromophobe (KICH), Kidney Renal Papillary Cell Carcinoma (KIRP), Lung Adenocarcinoma (LUAD), Lung Squamous Cell Carcinoma (LUSC), Stomach Adenocarcinoma (STAD), Thyroid Carcinoma (THCA), Uterine Carcinosarcoma (UCS) and Uterine Corpus Endometrial Carcinoma (UCEC).

The second most common and influential pathways are Axon guidance and Ras signaling pathways (tie) (Table 2). The Ras signaling pathway is in the top 10 most dysregulated pathways list for the following cancers: Lung Adenocarcinoma (LUAD), Lung Squamous Cell Carcinoma (LUSC), Thyroid Carcinoma (THCA), Breast Invasive Carcinoma (BRCA). The Axon guidance pathway is in the top 10 most dysregulated pathways list for the following cancers: Esophageal Carcinoma (ESCA) and Bladder Urothelial Carcinoma (BLCA).

**Table 2.**
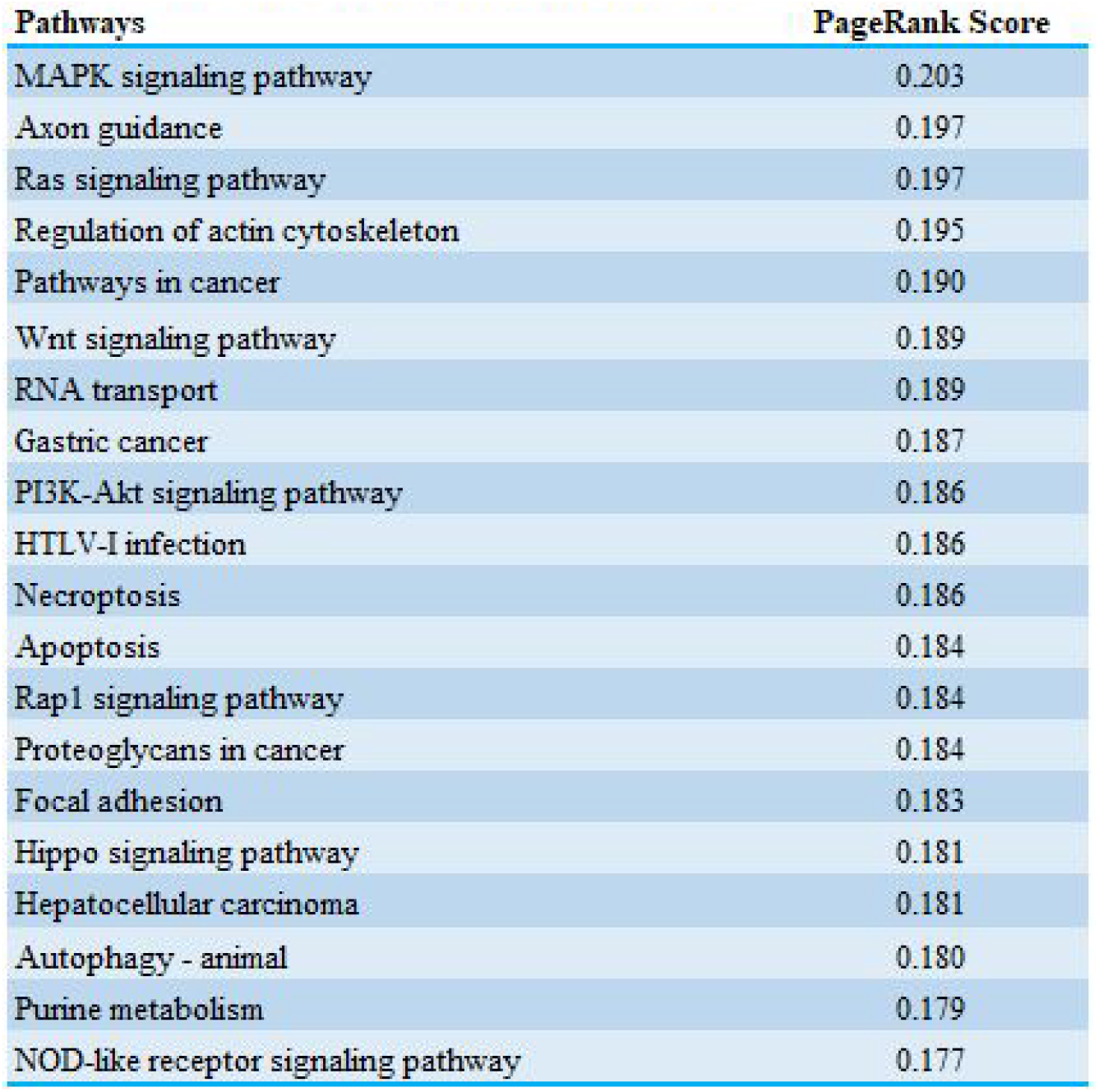
PageRank score of the pathways in the pan-cancer graph network.

| Pathways | PageRank Score |
| --- | --- |
| MAPK signaling pathway | 0.203 |
| Axon guidance | 0.197 |
| Ras signaling pathway | 0.197 |
| Regulation of actin cytoskeleton | 0.195 |
| Pathways in cancer | 0.190 |
| Wnt signaling pathway | 0.189 |
| RNA transport | 0.189 |
| Gastric cancer | 0.187 |
| PI3K-Akt signaling pathway | 0.186 |
| HTLV-I infection | 0.186 |
| Necroptosis | 0.186 |
| Apoptosis | 0.184 |
| Rap1 signaling pathway | 0.184 |
| Proteoglycans in cancer | 0.184 |
| Focal adhesion | 0.183 |
| Hippo signaling pathway | 0.181 |
| Hepatocellular carcinoma | 0.181 |
| Autophagy - animal | 0.180 |
| Purine metabolism | 0.179 |
| NOD-like receptor signaling pathway | 0.177 |

The third most common and influential pathway is Regulation of actin cytoskeleton (Table 2). The Regulation of actin cytoskeleton pathway is in the top 10 most dysregulated pathways list for the following cancers: Head and Neck Squamous Cell Carcinoma (HNSC) and Uterine Carcinosarcoma (UCS) and Uterine Corpus Endometrial Carcinoma (UCEC).

### Dysregulated Signaling Pathways

In the present study, the PoTRA algorithm detected the most dysregulated signaling pathways (by rank) in the pan-cancer analysis and those are the MAPK and PI3K-Akt signaling pathway. In the literature, the RAS-MAPK signaling pathways are the most critical in cancer because of the roles they have in relevant cellular processes such as survival, differentiation and proliferation (Masliah-Planchon, Garinet and Pasmant, 2015). Interestingly, the genes in the PI3K-Akt signaling pathway are highly mutated in cancer, and when dysregulated is found to support tumorigenesis, drug resistance and cancer progression (Mayer and Arteaga, 2016). Therefore, the frequent presence of the MAPK and PI3K-Akt signaling pathways (Table 3) in the list of dysregulated pathways detected by PoTRA, correlates with what is currently known in cancer.

**Table 3.** The significantly dysregulated signaling pathways in multiple cancers types.

|  |  |  |
| --- | --- | --- |
| **Cancer Projects** | **Signaling Pathways** | **Average Rank** |
| Thyroid Carcinoma (THCA) | PI3K-Akt Signaling Pathway | 1 |
| Breast Invasive Cancer (BRCA) | MAPK Signaling Pathway | 2 |
| Kidney Renal Clear Cell Carcinoma, Kidney Chromophobe, Kidney Renal Papillary Cell Carcinoma (KIRC, KICH, KIRP) | PI3K-Akt Signaling Pathway | 2 |
| Liver Hepatocellular Carcinoma (LIHC) | PI3K-Akt Signaling Pathway | 3 |
| Lung Adenocarcinoma and Lung Squamous Cell Carcinoma (LUAD, LUSC) | PI3K-Akt Signaling Pathway | 3 |
| Head and Neck Squamous Cell Carcinoma (HNSC) | MAPK Signaling Pathway | 4 |
| Colon and Rectum Carcinoma (COAD, READ) | MAPK Signaling Pathway | 5 |
| Prostate Adenocarcinoma (PRAD) | PI3K-Akt Signaling Pathway | 6 |
| Stomach Adenocarcinoma (STAD) | Rap1 Signaling pathway | 6 |
| Uterine Carcinosarcoma and Uterine Corpus Endometrial Carcinoma (UCS, UCEC) | MAPK Signaling Pathway | 6 |
| Esophageal Carcinoma (ESCA) | MAPK Signaling Pathway | 10 |
| Bladder Urothelial Carcinoma (BLCA) | Phosphatidylinositol Signaling | 16 |
| Glioblastoma Multiforme and Brain Lower Grade Glioma (GBM, LGG) | MAPK Signaling Pathway | 31 |
| Pancreatic Adenocarcinoma (PAAD) | Ras Signaling Pathway | 31 |
| Cholangiocarcinoma (CHOL) | TNF Signaling Pathway | 34 |
| Cervical Squamous Cell Carcinoma and Endocervical Adenocarcinoma (CESC) | Hedgehog Signaling Pathway | 35 |
| Adrenocortical Carcinoma, Pheochromocytoma and Paraganglioma (ACC, PCPG) | Phosphatidylinositol Signaling | 43 |

The PI3K-Akt signaling pathway is among the most dysregulated pathways in cancer, previous studies have reported somatic mutations in this pathway for liver, lung, prostate, thyroid cancers and an adrenal gland cancer subtype known as paraganglioma (Samuels and Velculescu, 2004; Grozinsky-Glasberg et al., 2008; Grozinsky-Glasberg et al., 2010; Harthill et al., 2002). The PoTRA algorithm also detected this highly ranked dysregulated pathway in the aforementioned cancer types (Table 3).

In a recent cancer study (Sikdar, Datta and Datta, 2016), a differential network analysis identified target pathways in several cancer sets from the International Cancer Genome Consortium (ICGC). The ICGC projects included Head and Neck Squamous Cell Carcinoma (HNSC), Lung Adenocarcinoma (LUAD) and Kidney Renal Clear Cell Carcinoma (KIRC). Overall, the study concluded that the PI3K-Akt and Ras signaling pathways are the most critical for HNSC, LUAD and KIRC (Sikdar, Datta and Datta, 2016). These results agree with the pan-cancer results (Table 3), with the exception of the TCGA HNSC project where the MAPK signaling pathway is the most dysregulated for this project. However, Ras signaling pathway does include the MAPK signaling pathway in it’s collection of pathways as seen in the KEGG DB (Kanehisa, Furumichi & Tanabe et al., 2017; Kanehisa, Sato & Kawashima et al., 2016; Kanehisa & Goto, 2000).

In the present study, the PoTRA algorithm identified the Rap1 signaling pathway as the most dysregulated signaling pathway for the stomach adenocarcinoma (STAD) project. In a recent TCGA study, the KIT gene was identified as a biomarker for STAD and was also found to activate the Rap1 signaling pathway (Pan et al., 2017). In the literature, the Rap1 signaling pathway is a key regulator of the multi-step process known as tumorigenesis, where the 3 major steps includes tumor cell migration, invasion, and metastasis (Zhang et al., 2017).

Other cancers such as BLCA and CHOL, are associated with cancer pathways in the literature such as PI3K-Akt signaling (Zheng et al., 2017) and MAPK as well as PI3K-Akt signaling (Rizvi et al., 2015) respectively. In the present study, the PoTRA algorithm identified dysregulated pathways such as Phosphatidylinositol signaling for BLCA and TNF signaling for CHOL that contain the reported pathways in their KEGG pathway networks (Kanehisa, Furumichi & Tanabe et al., 2017; Kanehisa, Sato & Kawashima et al., 2016; Kanehisa & Goto, 2000).

In recent studies, the cancers CESC (Chen et al., 2016) and PAAD (Logsdon and Lu, 2016) are both strongly associated with the dysregulated pathways identified by the PoTRA algorithm in this work, Hedgehog and Ras signaling pathways, respectively.

For the ACC (Maira et al., 2008) and PCPG cancers, the PI3K-Akt signaling pathway is a therapeutic target because it supports the proliferation of adrenocortical cancer cells (Zhikrivetskaya et al., 2017). The PoTRA algorithm also identified these pathways as the most dysregulated for these combined subtypes of adrenal gland cancer.

### Dysregulated Metabolic Pathways

The purine metabolism pathway is the most significantly dysregulated metabolic pathway (by rank) in 8 out of the 17 cancer types (Table 4). Although the purine metabolic pathway supports cellular proliferation and it is currently a therapeutic target for cancers, it’s biochemical regulators are still not fully understood (Pedley and Benkovic, 2017). However, the study of these biochemical mechanisms and how purine metabolism is dysregulated has increased the potential for new therapeutic approaches (Pedley and Benkovic, 2017). Although the purine metabolism pathway has been a therapeutic target for several years, many of the current therapeutics suffer from toxicity and thus motivates the study of purine metabolism pathway regulation (Pedley and Benkovic, 2017).

**Table 4.** Significantly dysregulated metabolic pathways in cancer.

| **Table 4 Significantly dysregulated metabolic pathways in cancer.** |  |  |
| --- | --- | --- |
| **Cancer Project** | **Metabolic Pathways** | **Average Rank** |
| Uterine Carcinosarcoma and Uterine Corpus Endometrial Carcinoma (UCS, UCEC) | Purine Metabolism | 8 |
| Stomach Adenocarcinoma (STAD) | Purine Metabolism | 10 |
| Liver Hepatocellular Carcinoma (LIHC) | Purine Metabolism | 10 |
| Head and Neck Squamous Cell Carcinoma (HNSC) | Purine Metabolism | 10 |
| Lung Adenocarcinoma and Lung Squamous Cell Carcinoma (LUAD, LUSC) | Purine Metabolism | 12 |
| Prostate Adenocarcinoma (PRAD) | Purine Metabolism | 20 |
| Bladder Urothelial Carcinoma (BLCA) | Inositol Phosphate Metabolism | 20 |
| Colon and Rectum Carcinoma (COAD, READ) | Glycerophospholipid Metabolism | 22 |
| Thyroid Carcinoma (THCA) | Pyrimidine Metabolism | 25 |
| Adrenocortical Carcinoma, Pheochromocytoma and Paraganglioma (ACC, PCPG) | Inositol Phosphate Metabolism | 30 |
| Kidney Renal Clear Cell Carcinoma, Kidney Chromophobe, Kidney Renal Papillary Cell Carcinoma (KIRC, KICH, KIRP) | Metabolism of Xenobiotics by Cytochrome P450 | 35 |
| Breast Invasive Cancer (BRCA) | Glycerophospholipid Metabolism | 40 |
| Pancreatic Adenocarcinoma (PAAD) | Glutathione Metabolism | 47 |
| Esophageal Carcinoma (ESCA) | Purine Metabolism | 50 |
| Cholangiocarcinoma (CHOL) | N-Glycan Biosynthesis | 50 |
| Glioblastoma Multiforme and Brain Lower Grade Glioma (GBM, LGG) | Purine Metabolism | 80 |
| Cervical Squamous Cell Carcinoma and Endocervical Adenocarcinoma (CESC) | Sphingolipid Metabolism | 82 |

A recent study of dysregulated metabolic pathways in BLCA, identified the Inositol phosphate metabolism pathway as one of the most dysregulated (Rodrigues et al., 2016). The PoTRA algorithm has also identified the Inositol phosphate metabolism pathway as one of the most dysregulated metabolic pathways for BLCA (Table 4).

For the COAD and READ cancers, the dysregulation of the Glycerophospholipid metabolism pathway is associated with cancer cell proliferation and tumorigenesis (Yan et al., 2016). This metabolic pathway was also identified by the PoTRA algorithm to be the most dysregulated pathway for the COAD and READ cancers (Table 4).

In the literature, the THCA cancer cells ensure their survival and proliferation by increasing their rate of glutamate metabolism to support their increasing need for glutamine (Guimaraes Coelho, Fortunato and Caravalho, 2018). Glutamate metabolism is part of the Pyrimidine metabolism network (Kanehisa, Furumichi & Tanabe et al., 2017; Kanehisa, Sato & Kawashima et al., 2016; Kanehisa & Goto, 2000) and involves the synthesis of purines and pyrimidines (Guimaraes Coelho, Fortunato and Caravalho, 2018). The PoTRA algorithm identified the Pyrimidine metabolism pathway as the most highly ranked dysregulated pathway for the THCA cancer type (Table 4).

In another study, ACC tumors increased their production of inositol phosphate as a result of an interaction between an AT1 receptor and phosphoinositidase C in NCI-H295R human cell lines (Parmar, Kulharya and Rainey, 2010). This finding correlates with PoTRAs identification of Inositol phosphate metabolism as the most dysregulated metabolic pathway for the ACC cancer.

For cancers such as PCPG, the mechanisms associated with the aggressiveness of this cancer are still not fully understood (Fliedner et al., 2012). However, a previous study found that PCPG patients with succinate dehydrogenase gene (SDHx) mutations have increased levels of inositol polyphosphate phosphatase 1 in their aggressive cancer cells as well as primary high-grade tumors (Fliedner et al., 2012). Additionally, a recent metabolites study found significantly increased levels of myoinositol in patients that were found to have SDHx mutations prior to developing PCPG (Imperiale et al., 2015). Therefore, PoTRAs identification of the inositol phosphate metabolism pathway as the most dysregulated pathway for PCPG has biological support.

Kidney renal cell carcinoma has three subtypes known as KIRC, KICH and KIRP (Li et al., 2018). In a recent study, the xenobiotic metabolism pathway was dysregulated in all 4 stages of the KIRC cancer (Li et al., 2018). Previous studies have associated the dysregulation of the xenobiotic metabolism pathway with drug resistance in KIRC cancer patients (Mitsui et al., 2015; Narjoz et al., 2014). Another recent study, found that the TCGA KIRC and KIRP project gene expression data that the xenobiotic metabolism pathways are indeed enriched (Schaeffeler et al., 2018). However, in the study, the enrichment score plot for the TCGA KICH project does not list xenobiotic metabolism as an enriched pathway (Schaeffeler et al., 2018).

The PoTRA algorithm identified the metabolism of xenobiotics by cytochrome P450 pathway as the most dysregulated pathway for the combined KIRC, KICH and KIRP cancers (Table 4). In the literature, cytochrome P450 (CYP) is a drug metabolizing enzyme and is expressed in the kidneys and other tissues (Molina-Ortiz et al., 2014). In experiments, the CYP enzyme has been found to have important roles in solid tumor chemoresistance and has also been found to interact (activate/inactivate) with chemotherapies (McFadyen, Melvin and Murray, 2004; Molina-Ortiz et al., 2014).

A recent BRCA study performed an integrated (proteomic and transcriptomic) enrichment analysis to determine if 228 untreated and primary BRCA tumor samples could be clustered into three metabolic groups (Mc1, Mc2 and Mc3) (Haukaas et al., 2016). The glycerophospholipid metabolic pathway is the most significantly different pathway between the subtypes Mc1 vs Mc3 and among the most significantly different pathways between Mc1 vs Mc2 (Haukaas et al., 2016). Therefore the glycerophospholipid metabolic pathway can be used to distinguish BRCA tumor subtypes (Haukaas et al., 2016). In the present study, the glycerophospholipid metabolic pathway was identified as the most dysregulated pathway for the TCGA BRCA project.

In a recent PAAD cancer study (Halbrook and Lyssiotis, 2017) proliferating cells used glutamine as the primary source of nitrogen and carbon, it was also found to maintain redox balance (Lyssiotis et al., 2013; Son et al., 2013). In redox balance, glutamine derived glutamate is utilized for glutathione synthesis (Halbrook and Lyssiotis, 2017). Another study identified glutathione as having a major role in cellular redox balance (DeBerardinis et al., 2007). Thus, the identification of the glutathione metabolism pathways as the most dysregulated pathway for the TCGA PAAD project has biological support.

In the literature, the ESCA, GBM and LGG cancers have been reported to have dysregulated purine metabolic pathways (Zhu et al., 2017; Strickland and Stoll, 2017; Wang et al., 2017). This agrees with the PoTRA results where purine metabolic pathway is identified as the most dysregulated pathway for the ESCA, GBM and LGG cancers (Table 4).

The CHOL cancer has been reported to have the N-glycan dysregulated pathway (Varki et al, 2017). The biosynthesis of N-glycans is upregulated to meet the metabolic needs of the cancer cells, therefore it’s inhibition will result in the prevention of further tumor growth (Varki et al, 2017). Thus, supporting PoTRAs identification of the N-Glycan pathway as the most dysregulated pathway for the CHOL cancer (Table 4).

In a recent study the sphingolipid metabolic pathway was reported as dysregulated in the CESC cancer (Porcari et al., 2018). The dysregulation of the sphingolipid metabolic pathway may induce cellular processes known as apoptosis and cellular autophagy (Lizardo et al., 2017). This finding supports PoTRA’s identification of the sphingolipid metabolic pathway as the most dysregulated pathway in the CESC cancer (Table 4).

### Most Highly Ranked Dysregulated Pathways

Pathways in cancer is the most common dysregulated pathway in 8 of 17 cancer types in the pan-cancer analysis, ranking in the top 10 (Table 5). Pathways in cancer encompasses a set of pathways often dysregulated in cancer, including MAPK, PI3K-Akt, TGFB and Jak-Stat signaling, etc (Kanehisa, Furumichi & Tanabe et al., 2017; Kanehisa, Sato & Kawashima et al., 2016; Kanehisa & Goto, 2000). Therefore, it’s not surprising that multiple cancer projects were found to be associated with the pathway known as Pathways in cancer.

**Table 5.** Significantly dysregulated pathways in cancer.

| **Table 5 Significantly dysregulated pathways in cancer.** |  |  |
| --- | --- | --- |
| **Cancer Project** | **Pathways** | **Average Rank** |
| Breast Invasive Cancer (BRCA) | Pathways in Cancer | 1 |
| Head and Neck Squamous Cell Carcinoma (HNSC) | Pathways in Cancer | 1 |
| Lung Adenocarcinoma and Lung Squamous Cell Carcinoma (LUAD, LUSC) | Pathways in Cancer | 1 |
| Stomach Adenocarcinoma (STAD) | Pathways in Cancer | 1 |
| Thyroid Carcinoma (THCA) | PI3K-Akt Signaling Pathway | 1 |
| Colon and Rectum Carcinoma (COAD, READ) | Pathways in Cancer | 2 |
| Kidney Renal Clear Cell Carcinoma, Kidney Chromophobe, Kidney Renal Papillary Cell Carcinoma (KIRC, KICH, KIRP) | Pathways in Cancer | 2 |
| Liver Hepatocellular Carcinoma (LIHC) | Pathways in Cancer | 2 |
| Prostate Adenocarcinoma (PRAD) | Fluid Shear Stress and Atherosclerosis | 2 |
| Bladder Urothelial Carcinoma (BLCA) | Autophagy - Animal | 3 |
| Uterine Carcinosarcoma and Uterine Corpus Endometrial Carcinoma (UCS, UCEC) | Pathways in Cancer | 8 |
| Cholangiocarcinoma (CHOL) | Vibrio Cholerae Infection | 9 |
| Cervical Squamous Cell Carcinoma and Endocervical Adenocarcinoma (CESC) | Necroptosis | 13 |
| Adrenocortical Carcinoma, Pheochromocytoma and Paraganglioma (ACC, PCPG) | Inositol Phosphate Metabolism | 30 |
| Glioblastoma Multiforme and Brain Lower Grade Glioma (GBM, LGG) | MAPK Signaling Pathway | 31 |
| Pancreatic Adenocarcinoma (PAAD) | Ras Signaling Pathway | 31 |
| Esophageal Carcinoma (ESCA) | Purine Metabolism | 50 |

In the literature, the THCA cancer is associated with a dysregulated PI3K-Akt signaling pathway (Jin et al., 2013). The dysregulated PI3K-Akt signaling pathway supports the survival response in THCA cancer cells, therefore this pathway has served as a therapeutic target multiple times (Jin et al., 2013). This supports PoTRAs identification of this pathway as the most dysregulated for the THCA project (Table 5).

In two recent studies, low fluid shear stress was found to induce tumor cell metastasis in cancer (Huang et al., 2018) and in at least one study, tumor metastasis was driven by the stimulation of the YAP1 gene, in the PRAD cancer (Lee et al., 2017). Thus supporting PoTRA’s conclusion that the fluid shear stress and atherosclerosis pathway is the most dysregulated pathway for the TCGA PRAD project (Table 5).

In the BLCA cancer the Autophagy - animal pathway is dysregulated to support the survival of cancer cells during metabolic stress (Chandrasekar and Evans, 2016; Chen and Karantza, 2011; Leone and Amaravadi, 2013). Therefore, PoTRA’s identification of the Autophagy - animal pathway as the most dysregulated pathway in the TCGA BLCA project (Table 5).

In a recent TCGA study, oxidative phosphorylation is altered and targeted for inhibition in CHOL cancer (Farshidfar et al., 2017). Thus supporting PoTRA’s identification of the Vibrio Cholerae Infection pathway as the most dysregulated pathway in the TCGA CHOL project, because oxidative phosphorylation is part of the Vibrio Cholerae Infection pathway network (Table 5).

Many cancer cells have a dysregulated Necroptosis pathway, this includes CESC cancer cells (Su et al., 2016). Cancer cells eliminate necroptosis mechanisms and/or develop resistance against them (Su et al., 2016). This supports PoTRA’s conclusion that the Necroptosis pathway is the most dysregulated pathway in the TCGA CESC project (Table 5).

The dysregulated pathways Inositol phosphate metabolism and Purine metabolism for the ACC and PCPG as well as ESCA cancers were discussed in the Dysregulated Metabolic Pathways section. The dysregulated MAPK and Ras signaling pathways for the GBM and LGG as well as PAAD cancers were discussed in the Dysregulated Signaling Pathways section.

## DISCUSSION

Carcinogenesis is induced by a combination of damaging environmental stresses that includes exposures to carcinogens, UV radiation and heat shock, (NIH-NCI 2015; Sonenberg and Hinnebusch 2009) and intracellular cues such as the dysregulation of key pathway genes (Wilk and Braun, 2018). The purpose of the PoTRA algorithm is to use Google’s PageRank algorithm to find the hub genes in normal and tumor pathway networks, and to thus determine if there is truly a difference between these networks (i.e., determine if the pathway is dysregulated).

Overall, the PoTRA tool can accurately identify dysregulated pathways even in tumor samples with intratumoral heterogeneity by using a majority-rules approach. Interestingly, tumors such as those from the TCGA projects ACC, PCPG and GBM, LGG have different gene expression profiles (as seen in PCA plots) yet are still associated with the same dysregulated pathways respectively.

Also, the PoTRA algorithm can identify dysregulated pathways by using only protein-coding mRNA while other algorithms arrive at similar conclusions by using a multi-omics approach. The robustness of the PoTRA algorithm (i.e., the least average standard deviation in the ranks of the detected mRNA dysregulated pathways) can be guaranteed only if the dataset is balanced and has a minimum of 50 samples per phenotype. Thus, the most consistent ranks were found for the following primary sites: Breast, Colorectal, Kidney, Liver, Lung, Prostate and Thyroid.

## FUTURE DIRECTION

The PoTRA algorithm will continue to be developed so that it can utilize a more comprehensive and integrated approach to determine how that can impact the accuracy of dysregulated pathway identification. Additionally, visualizations for the graph networks will also be further developed.

## CONCLUSION

The PoTRA algorithm accurately identified the dysregulated pathways associated with each TCGA cancer project. It was also easily used in the cancer genomics cloud and therefore is suited for large scale analyses.

## Acknowledgments

Thank you to the technical support team at the Cancer Genomics Cloud. The Seven Bridges Cancer Genomics Cloud has been funded in whole or in part with Federal funds from the National Cancer Institute, National Institutes of Health, Contract No. HHSN261201400008C and ID/IQ Agreement No. 17X146 under Contract No. HHSN261201500003I

